# Development and Characterization of Self-Tracing Neural Progenitor Cells for Mapping Their Synaptic Integration into Endogenous Neural Networks

**DOI:** 10.64898/2026.02.20.706998

**Authors:** Priscilla Chan, Zijian Lou, Christopher S Ahuja, Dylan Gauthier, Alexander A. Velumian, Mohamad Khazaei, Michael G. Fehlings

## Abstract

Neural progenitor cell (NPC) transplantation holds immense promise for neurodegenerative and traumatic central nervous system (CNS) pathologies. However, it is crucial to define which neural circuits and pathways are targeted with transplanted NPCs under different conditions. A major roadblock lies in the limited ability to accurately trace integration of grafted cells into the host neural network. Conventional tracers suffer from drawbacks like low trans-synaptic efficiency, toxicity, and difficulty in efficiently and specifically targeting transplanted cells.

To address these critical limitations, we have developed self-tracing NPCs genetically engineered to express both anterograde (WGA-mCherry) and retrograde (GFP-TTC) trans-synaptic tracers. These self-tracing NPCs maintain their intrinsic properties, differentiate into electrically active neurons, and integrate into host circuitry *in vitro*. Importantly, co-culture with primary rat neurons revealed successful trans-synaptic tracing of grafted human neurons, evidenced by single-positive WGA+ or TTC+ rat cells. *In vivo*, NPCs transplanted into a rodent spinal cord injury model retained tracer expression for 12 weeks, enabling visualization of grafted cells within the spinal cord. Co-labeling with WGA and TTC provided evidence that these NPCs forms neurons which integrated into local circuits. Our novel self-tracing NPC platform offers a powerful tool to overcome trans-synaptic tracing challenges. This approach provides the opportunity to gain critical insights into graft integration and neural circuit remodeling, paving the way for better-designed transplantation strategies and improved therapeutic outcomes in a broad spectrum of CNS disorders.

## Introduction

The intricate neural circuits of the central nervous system, once disrupted by injury or disease, present a profound challenge for repair and regeneration. Neural progenitor cells (NPCs), with their potential to replace lost neurons and bridge severed connections, offer promising treatment options to restore these lost connections. A pivotal aspect of employing NPCs is ensuring their effective integration into pre-existing neural networks, thereby re-establishing functional synapses essential for recovering impaired neurological functions^1,2^. However, a major obstacle in this domain has been the absence of precise methodologies to track and comprehend how NPCs incorporate into the host’s neural architecture.

Traditional tracing methods, which aim to reveal these emerging connections, are often limited by issues such as toxicity, weak signal intensity, and an inability to track complex, multi-synaptic integration processes. Non-viral tracers, like horseradish peroxidase and fluorogold, are useful in some contexts but are largely non-trans-synaptic. While some non-viral trans-synaptic tracers exist, such as cholera toxin B^3^, their inability to replicate results in signal dilution across the synapse, leading to potentially misleading interpretations of synaptic connectivity^4^. These limitations often necessitate the use of viral tracers, such as herpes simplex virus (HSV)^5^. and pseudorabies virus (PRV), which are preferred for trans-synaptic tracing due to their ability to replicate within host neurons and infect adjacent synaptic connections, thereby amplifying the signal^6–8^. However, the toxicity of viral techniques, although beneficial for their specificity and wide applicability, restricts their practical use to mainly monosynaptic tracing.

In this paper, we developed a novel approach for tracing transplanted cells by making self-tracing NPCs. These genetically modified cells are equipped with two potent tracers, one for anterograde tracing (mapping outgoing neural pathways) and one for retrograde tracing (mapping incoming neural pathways). The first tracer utilized is Wheat Germ Agglutinin (WGA), a plant lectin that binds specifically to neural membrane glycoproteins and glycolipids, enabling anterograde transport and labeling of neural pathways^9, 10, 11^. WGA has been fused to mCherry for visualization. The second tracer is Tetanus Toxin Fragment C (TTC) ^12, 13^, known for its retrograde axonal transport properties, fused to GFP^14, 15^. This combination of WGA and TTC, engineered into NPCs, provides a comprehensive method for visualizing the integration patterns of NPCs as they establish connections with the host’s neural circuitry. This approach not only overcomes the limitations of conventional methods but also offers a more direct and reliable method to study synaptic integration in vivo.

The ability of these NPCs to trace their own synaptic connections post-transplantation into the CNS offers a unique tool for understanding the complex dynamics of neural regeneration and integration. The use of WGA and TTC, each with its specific transport properties, enables detailed mapping of both anterograde and retrograde synaptic connections, providing critical insights into the integration process of grafted cells within the host neural network.

The findings from this study have the potential to significantly impact the field of neural stem cell regenerative therapeutics. By mapping the connectivity facilitated by self-tracing NPCs, new avenues for targeted and effective therapeutic strategies can be developed. This offers insights that could inform the design of future NPC-based therapies, aiming to not only replace lost neurons but also to seamlessly integrate into the CNS, thus restoring function. This exploration into the connectivity of self-tracing NPCs marks a step towards a future where severe neural disruptions might be effectively repaired.

## Results

### Expression of WGA-mCherry and GFP-TTC in NPCs

NPCs were transfected with bicistronic constructs expressing both WGA and TCC that were fused to mCherry and GFP, respectively. We integrated the expression cassette into the host genome using the piggyBac transposon (Figure 1A-D). Transfected NPCs were screened for mCherry and GFP expression as a proxy for WGA and TTC expression. Initially, approximately 1% of a 10 cm plate displayed dual-positive expression of the markers. After FACS, individual cells were expanded over 2 weeks before the second round of fluorescent screening was performed using an epifluorescence microscope. Only cells maintaining ubiquitous expression of both mCherry and TTC were passaged for further growth (Figure 1C, D).

**Figure 1:**
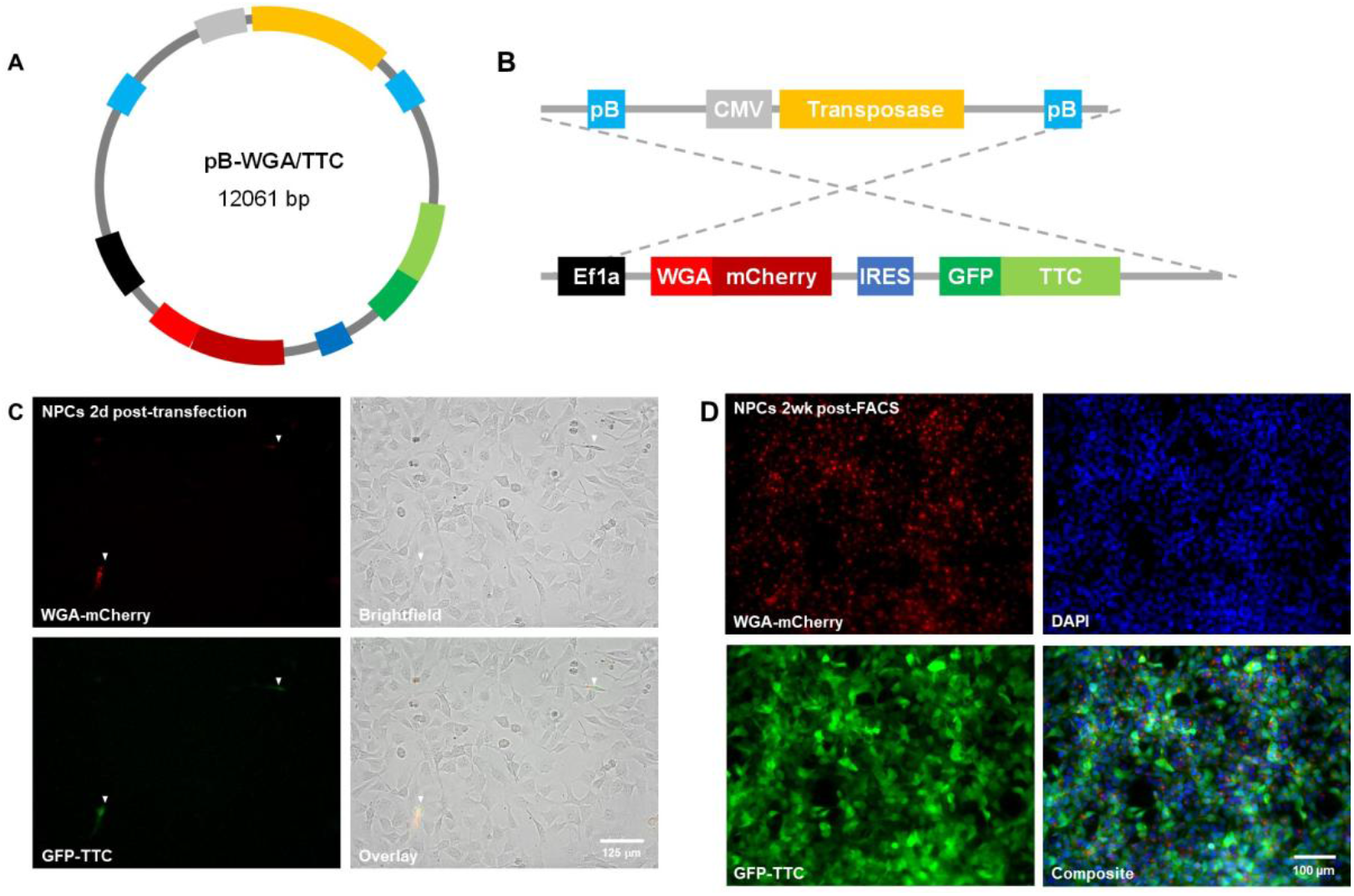
The plasmid map of self-tracing NPCs. (A) Plasmid Map of the Self-Tracing Construct in *Piggy Bac* Vector (B) Box-line diagram highlighting key components of the DNA sequence, including constitutive promoters CMV and EF1a, transposase enzyme for genome integration, fusion protein tracers WGA-mCherry (anterograde), and GFP-TTC (retrograde). (C, D) Monoclonal line of self-tracing NPCs. (C) Approximately 1% of NPCs demonstrated tracer expression 2 days post-transfection. (D) Ubiquitous expression of both WGA-mCherry and GFP-TTC was seen in NPCs 2 weeks post-FACS. Representative images are shown for WGA TTC.

### Self-tracing NPCs retain self-renewal and proliferative properties

WGA TTC self-tracing NPCs were tested for self-renewal and proliferative potential by neurosphere assay against a control line of GFP-expressing NPCs, previously generated and tested in our lab (Figure 2). The number of primary neurospheres greater than 50 μm was comparable between all NPC lines (GFP: 36.0 ± 2.8; WGA/TTC: 32.8 ± 3.0). Similar numbers of neurospheres were also observed after the first (GFP: 32.0 ± 1.8; WGA/TTC: 33.6 ± 2.0) and second passages (GFP: 37.6 ± 3.0; WGA/TTC: 35.2 ± 3.4). Self-tracing NPCs also maintained tracer expression (Figure 2B). Further characterization of NPC markers by RT-PCR has been included in Supplemental Figure 1.

**Figure 2:**
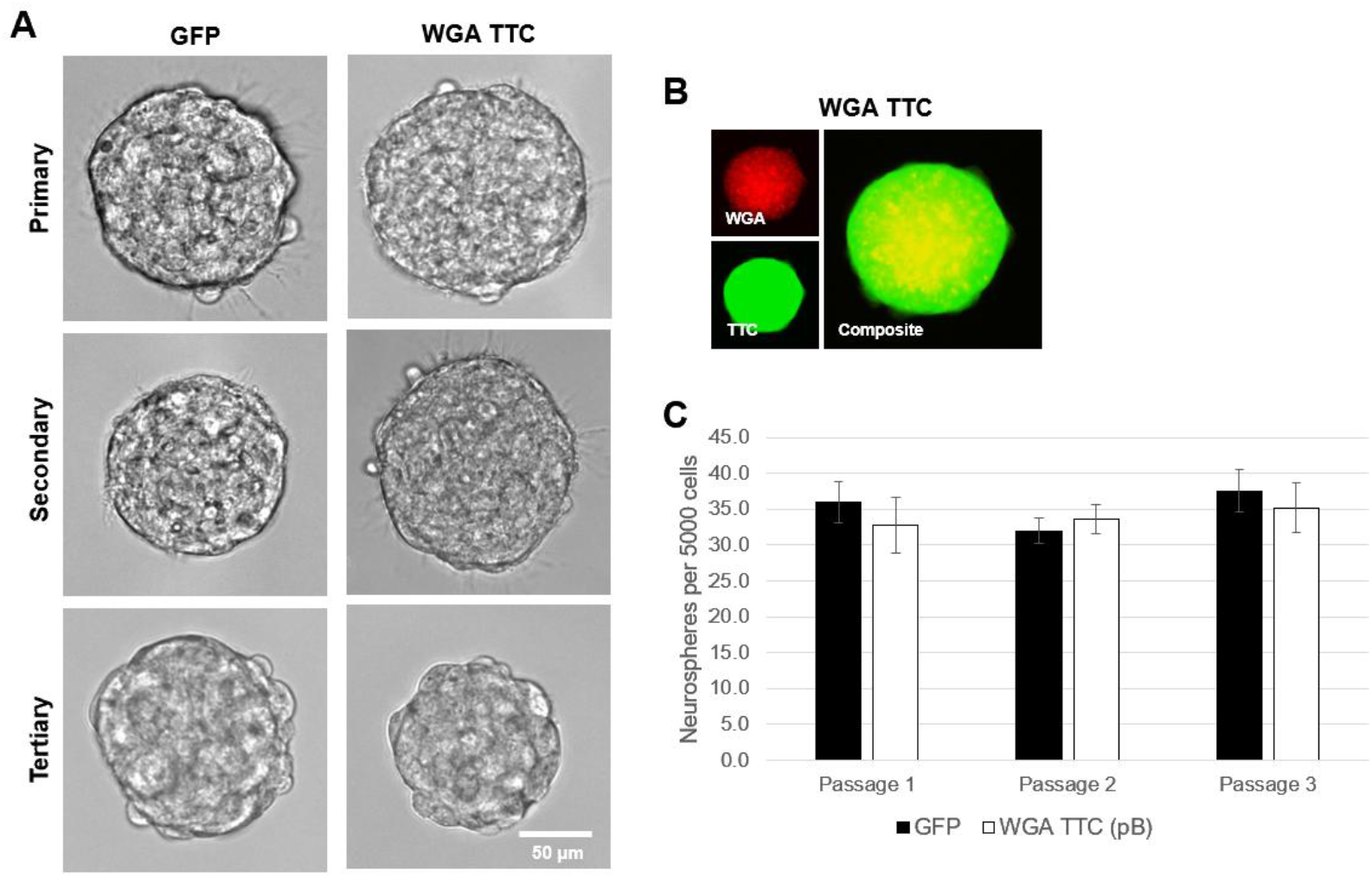
Self-tracing NPCs demonstrate typical proliferative properties. (A) GFP and self-tracing NPCs were cultured for 7 days to form primary neurospheres. Neurospheres were subsequently triturated and plated for an additional 7 days to form secondary and tertiary neurospheres. Representative brightfield images are shown for all cell lines. (B) Fluorescent images of both self-tracing lines of NPCs demonstrate ubiquitous expression of WGA and TTC. (C) After transferring and fixing neurospheres to a Geltrex-coated plate, the number of neurospheres greater than 50 μm was quantified. Genetically modified NPCs did not show any difference in proliferative or self-renewal properties. Error bars display SEM, n=3 per group. One-way ANOVA corrected for multiple comparisons by Tukey’s post hoc test, n.s. (p > 0.05). Abbreviations: GFP = green fluorescent protein; pB = piggyBac; TTC = tetanus toxin fragment C; WGA = wheat germ agglutinin.

### *In vitro* induction of NPC differentiation to neurons

NPCs need to be able to differentiate into neurons to replace lost neuronal connections and integrate into the host circuitry in disease conditions. To ensure the self-tracing NPC line could differentiate into neurons and form traceable connections, the cells were cultured in a cocktail of growth factors and stained for neuronal markers. After 1 week in culture with Neural Induction Media (NIM), self-tracing NPCs stained positive for the immature neuron marker TUBB3. Switching to Neural Maintenance Media (NMM) for an additional week in culture, self-tracing NPCs expressed MAP2, a cytoskeletal marker found in more mature neurons (Figure 3B). When plated within Campenot chambers, neurons could be seen growing axons up to 800 μm (Figure 3E). Self-tracing NPC differentiation to astrocytes and oligodendrocytes was also studied by *in vitro* differentiation. Self-tracing NPCs displayed robust GFAP expression, typical of astrocytes. Further characterization of glial differentiation by RT-PCR has been included in Supplemental Figure 2.

**Figure 3:**
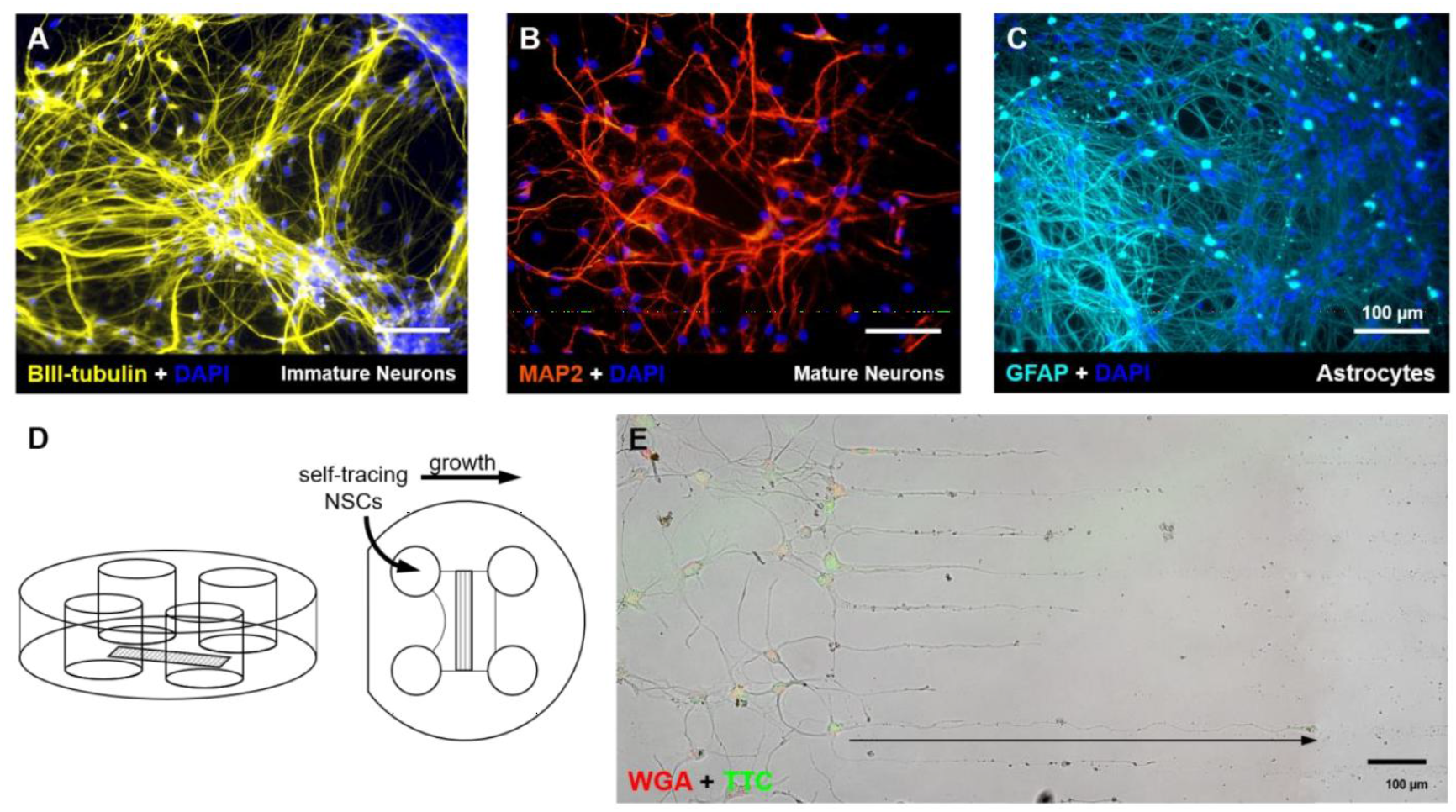
Self-tracing NPC can differentiate into neurons and astrocytes. (A, B) Self-tracing NPCs treated with neurobasal media with a cocktail of growth factors can differentiate into neurons. (A) Immature neurons were detectable within 1 week of induction and (B) more mature neurons were seen after 2 weeks. (C) Using 1% FBS, self-tracing NPCs became astrocytes within 1 week of induction. Representative images are shown of self-tracing cells. Images were taken at 20X. (D) Diagram of Campenot chamber used to see directed axon growth. (E) NPCs that differentiated into neurons over 2 weeks in a Campenot chamber were seen to grow long axons.

### Self-tracing NPC-derived neurons are electrophysiologically functional

After confirming that the self-tracing NPCs can differentiate into neurons, their ability to form electrically functional synapses was investigated. If the synapses formed by the NPCs are not electrically active, then the connections that grafted NPCs make with the host circuitry may be non-functional and confer no benefit to recovery. Furthermore, functional neurons are required not only for integration after transplantation but also for transmission of trans-synaptic tracers. Whole-cell patch clamp recording was used to evaluate the neuronal electrophysiological phenotype of self-tracing NPC-derived neurons. NPCs were differentiated for a total of 4 weeks prior to recording, in hopes of studying more mature neurons. Neurons were identified by dual-positive expression of mCherry and GFP under epifluorescence. Whole-cell voltage clamp recordings were used to probe for the presence of voltage-activated Na^+^ and K^+^ currents. As seen in Figure 4C, depolarizing voltage steps activated a fast inward current (the downward peak in Fig 4C) pointing to the presence of voltage-gated Na^+^ channels typical of neurons. The inward Na^+^ current was followed by a slower and more prolonged outward K^+^ current. The ability of the cells to generate action potentials was confirmed by depolarizing current steps in current clamp recording (Fig 4D). Together, these functional hallmarks of neuronal phenotype suggested that self-tracing neurons have the potential to integrate into host circuitry and trace their synaptic connections.

**Figure 4:**
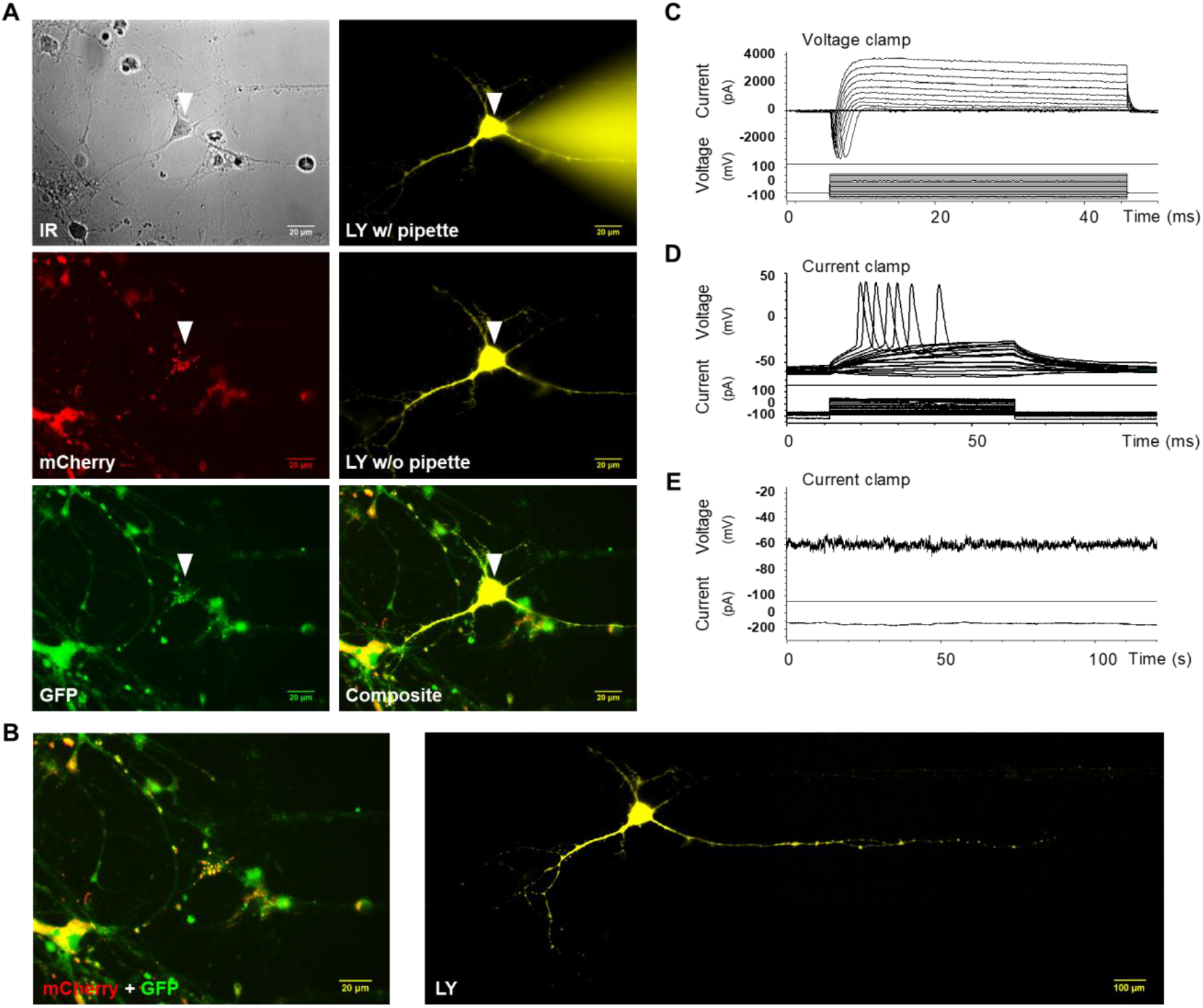
Self-tracing NPC-derived neurons are electrically functional. (A) Self-tracing NPCs were differentiated to neurons over 4 weeks. Cells were visualized using differential infrared contrast microscopy (IR in A) and epifluorescent microscopy to identify neurons for whole-cell recording. Arrowheads in A indicate the patched cell corresponding to the electrophysiological recordings shown in C-D. During whole-cell recording, the neurons were injected with Lucifer Yellow (LY) through the patch pipette. Representative images of WGA-TTC NPCs are shown. (B) Merged image of mCherry and GFP expression confirm the expression of both molecular tracers, WGA-mCherry and GFP-TTC, in patched cells. Stitched image of a LY-injected neuron shows the complete cell morphology. (C) Voltage clamp recording revealed a large inward Na+ current followed by an outward K+ current, typical of neurons. (D) Current clamp recording demonstrated that differentiated neurons fire action potentials once the membrane potential reached threshold. (E) Two min voltage recordings in current clamp mode did not reveal any spontaneous synaptic activity. The membrane potential of the cell was held at near -60 mV by injected current (shown on the lower trace).

### *In vitro* and *in vivo* analysis of self-tracing NPCs

Once the functional activity of self-tracing NPC-derived neurons was confirmed, activity dependent trans-synaptic tracing could be tested by co-culturing self-tracing neurons with primary rat neurons. This would serve as a proxy for lengthier transplantation of NPCs into an animal. Self-tracing NPCs underwent 2 weeks of neural induction prior to the addition of primary rat cortical neurons isolated from E17-19 pups. Co-culture was performed for an additional 2 weeks. Longer co-cultures were attempted but increased the risk of bacterial contamination and primary cell death. To differentiate between human and rat cells, STEM121 staining was used to reveal the distribution of human self-tracing neurons (Figure 5B). MAP2 staining further confirmed the presence of all neurons in culture, including rat primary neurons. MAP2 was paired with a quantum dot Q565 antibody to prevent emission spectrum overlap with other markers. However, Q565 excitation overlapped with DAPI and resulted in unusual nuclear staining. Nevertheless, WGA staining was clearly seen surrounding DAPI in STEM121-rat neurons (Figure 5C). Cytosolic TTC signal was also seen in a smaller population of rat neurons. Co-culture of wild-type NPCs, without any fluorescent tags, with rat primary neurons showed no WGA or TTC signal. Together, these findings demonstrate that differentiated self-tracing NPCs indeed integrate with rodent neurons and form synaptic connections.

**Figure 5:**
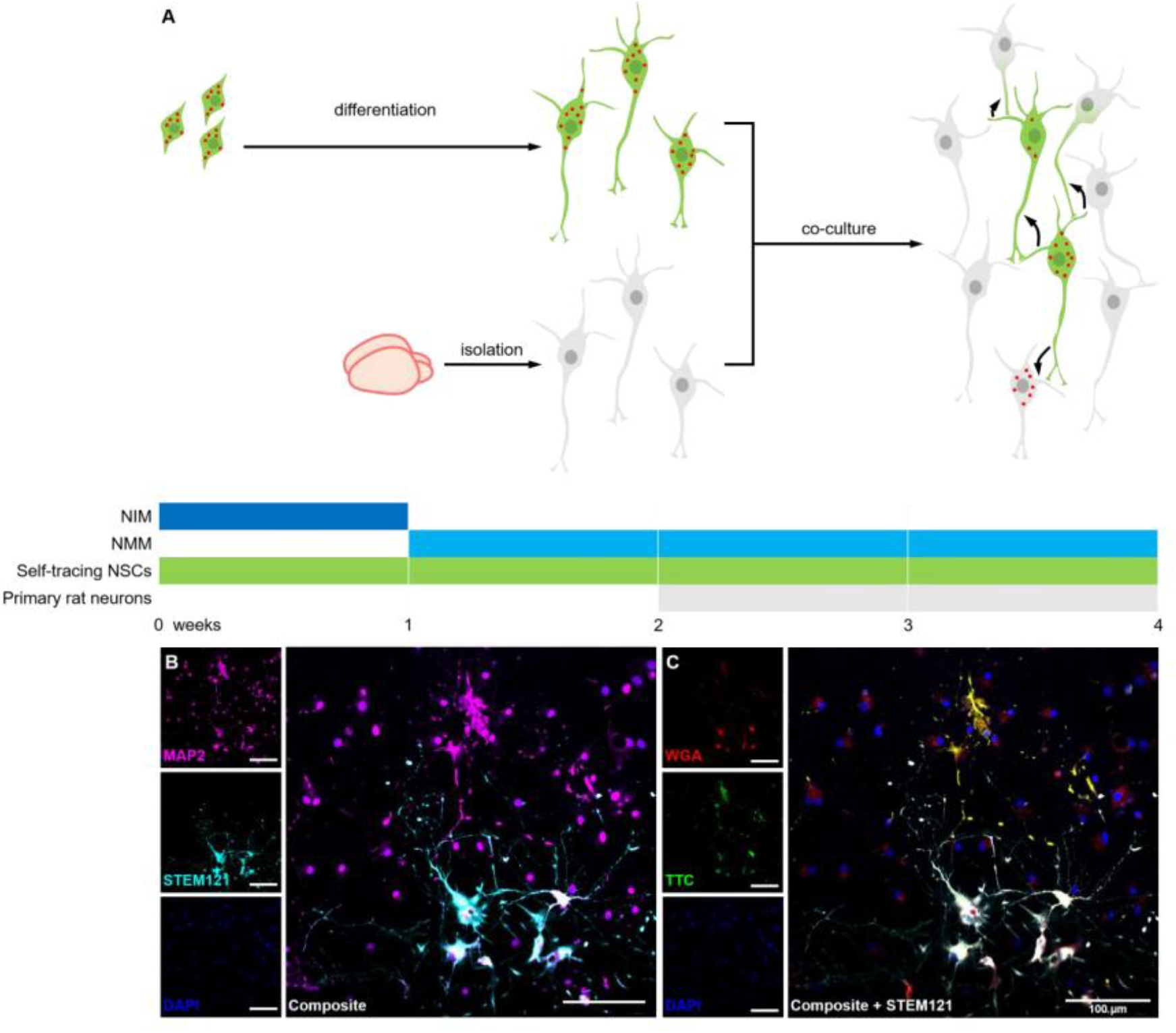
Self-tracing NPCs integrate and trace primary rat neurons *in vitro*. (A) Design of *in vitro* co-culture of self-tracing NPCs with primary rat cortical neurons. Self-tracing NPCs were plated and differentiated to neurons for 2 weeks prior to adding primary rat cortical neurons. Cells were left to mature and integrate over an additional 2 weeks. (B) Representative image of co-culture. STEM121 reveals distribution of human self-tracing NPC-derived neurons while MAP2 stains all neurons, including rat cells. (C) The same field of view shows dual positive WGA and TTC expression in human cells, but single WGA+ or TTC+ rat neurons.

### Self-tracing NPCs maintain tracer expression 12 weeks post-transplant

Moving towards transplantation in a rodent model of SCI, RNU rats were injured with a 23 g contusion-compression model at the C6/7 level. Eight weeks post-SCI, self-tracing NPCs were prepared for transplantation at the epicentre and 4 additional sites (1 mm rostral and caudal to the lesion at a depth of 1 mm, bilaterally). Animals were kept for 12 weeks post-transplantation to allow for migration, differentiation, and integration of grafted cells into the host tissue. Firstly, long-term transgene expression of WGA-mCherry and GFP-TTC was examined (Figure 6). IHC demonstrated sufficient WGA and TTC signal when enhanced with antibody staining. Secondly, to see whether tracing occurred, we focused on finding cells that were only single-positive WGA+ or TTC+ cells. Distinct single-positive cells were identified, suggesting that tracing occurred.

**Figure 6:**
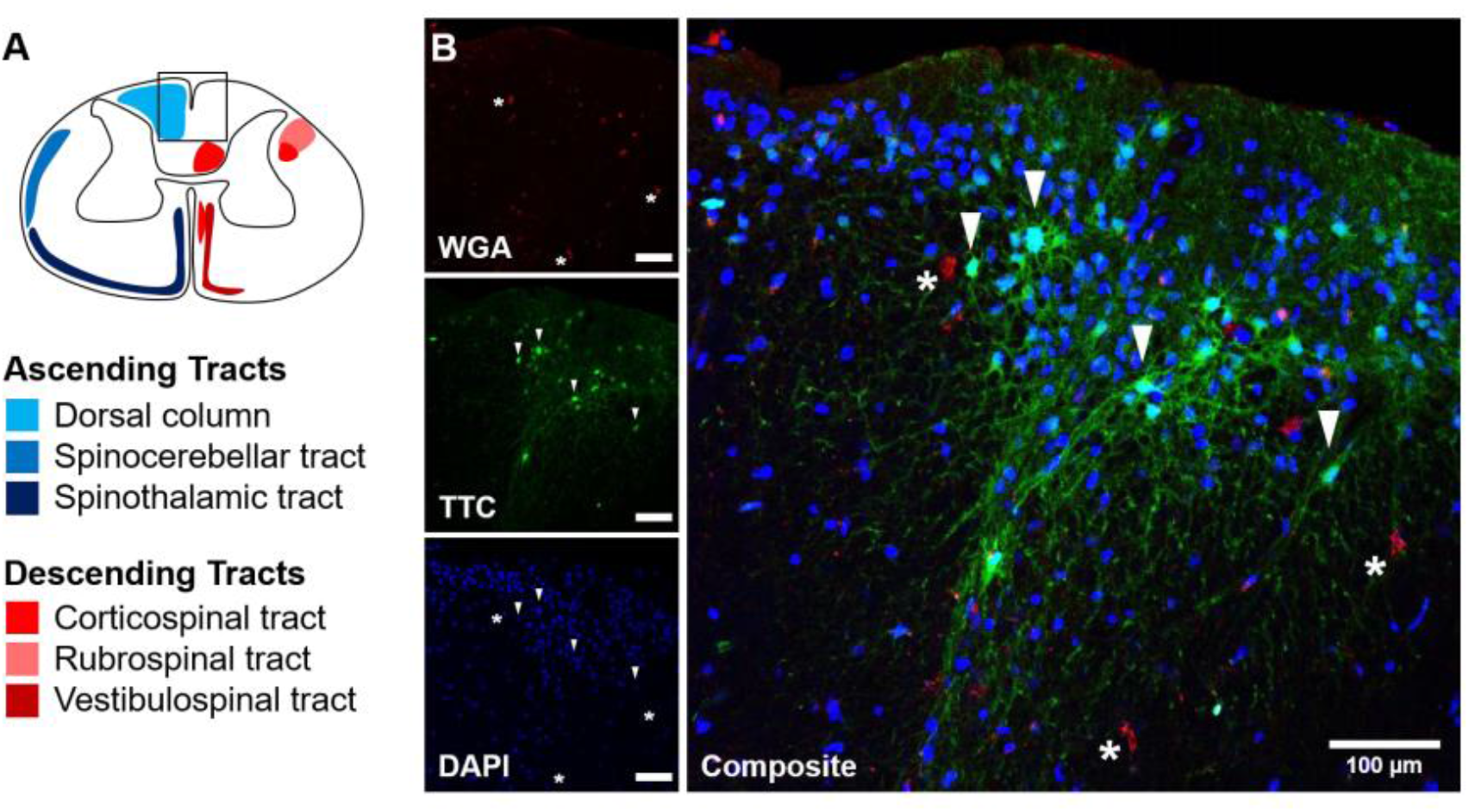
Self-tracing NPCs maintain tracer expression 12 weeks post-transplant. (A) Tract organization in a rodent cervical spinal cord for reference. The box marks the approximate region shown in the representative image. (B) Strong tracer expression is still observed 12 weeks post-transplantation. Asterisks mark WGA+/TTC-cells while arrowheads mark WGA-/TTC+ cells. Representative image of the dorsal funiculus at the epicentre of a 12 week-post self-tracing NPC transplant into an 8-week SCI RNU rat (23 g clip compression injury at the C6/7 level). Image taken at 20X.

## Discussion

This study marks a paradigm shift in neural tracing, specifically in the field of regenerative neural cell transplantation. We have successfully developed and characterized self-tracing NPCs, genetically engineered to express dual trans-synaptic tracers, WGA-mCherry and GFP-TTC^16^. This innovative approach addresses a critical gap in neurobiological research, moving beyond limitations and potential neurotoxicity associated with conventional viral tracers.

Unlike past studies largely focused on host neuron plasticity, our self-tracing NPCs offer a method to express tracers internally eliminates the need for injections and their inherent targeting difficulties. More importantly, it bypasses the biosafety concerns and complexities of viral vectors, a crucial advantage for therapies like SCI treatment, where precise synaptic integration mapping is vital.

Our co-culture experiments provide compelling evidence of successful neuron tracing. The presence of single-positive WGA and TTC neurons without STEM121 confirms the specific tracing of human neurons derived from self-tracing NPCs. These findings not only reaffirm the suitability of WGA and TTC as trans-synaptic tracers but also hint at the potential morphology of integrated neurons. Interestingly, the observed pyramidal-like morphology in Figures 5 and 6 suggests long axon projections and aligns with upper motor neuron characteristics (e.g., corticospinal tract)^17^. Future studies could explore this intriguing possibility after in vivo transplantation.

Twelve weeks post-NPC transplant, our in vivo study reveals GFP+ and mCherry+ cells in the dorsomedial aspect of the dorsal funiculus. While single-positive cells suggest potential tracing, definitive neuron identification requires co-labeling with neuronal markers and future anatomical investigations. Nonetheless, the observed distribution within the dorsal white matter, close to the dorsal column medial lemniscus (DCML) location, warrants further investigation. This raises exciting possibilities for both regeneration and relay through DCML and CST fibers.

This study establishes the feasibility of genetically modifying hiPSC-NPCs to express tracers while maintaining their crucial characteristics. The sustained tracer expression up to 12 weeks highlights the stability of our approach and opens doors for chronic studies of neural connectivity. While future research needs to address grafted cell identification limitations, our self-tracing NPCs offer a powerful tool for studying NPC integration and optimizing transplantation strategies. Importantly, the successful in vitro tracing and potential in vivo tracing demonstrate the immense potential of this technology to guide the development of effective, targeted therapies for a range of CNS disorders.

While NPCs have advanced to clinical trials for treating SCI, the translation of significant neurological recovery from preclinical studies to patient populations has been limited. A crucial aspect missing in the advancement of stem cell therapies is the understanding of specific motor and sensory pathways that contribute to recovery. Essential to this understanding is the development of improved methods for tracing the trans-synaptic integration of grafts within host circuitry. Our research makes a pivotal contribution in this area by introducing NPCs capable of mapping synaptic connections following transplantation. The implications of our self-tracing NPCs extend well beyond SCI treatment. Considering the versatility of NPCs and the fundamental role of synaptic connectivity in the CNS, these cells hold immense potential for investigating a range of neurodegenerative and traumatic conditions.

## Material and Methods

### Plasmid subcloning

The fusion protein WGA-mCherry was inserted in piggyBac vector^18^ under an EF1a promoter by Gibson assembly, and GFP-TTC was subcloned downstream of WGA-mCherry under the IRES to generate a bicistronic vector (Figure 1).

### Generation of the cell line

NPCs were previously generated through the dual SMAD inhibition protocol from a hiPSC line ^18^. The resultant hiPSC-NPCs were maintained in culture in N2/B27 Serum Free Media (SFM). Cells were transfected with the self-tracing construct through electroporation using Amaxa Nucleofector. To reduce cell death, SFM with Rho/ROCK inhibitor (10 μM, Y-27632) was added to the cuvette and cells were gently transferred to a new Geltrex-coated plate. Expression of mCherry and GFP was monitored over the course of 2 weeks. Cells then underwent fluorescent activated cell sorting (FACS) into coated 96-well plates. SFM change was done the following day to remove the Rho/ROCK inhibitor and Ca2+ from the sheath fluid. Subsequent media changes were done once a week until fluorescence could be visualized. Two weeks post-FACS, NPCs were screened for GFP and mCherry expression, with only strong dual-positive cells being passaged for expansion.

### Neurosphere assay

To assess self-renewal and proliferative potential, self-tracing NPCs and control GFP-NPCs were plated at a clonal density of 10 cell/μL in SFM on uncoated 24-well suspension plates. Neurospheres grew undisturbed for 7 days. Neurospheres greater than 50 μm in diameter were quantified. Just prior to imaging, the contents of each well were transferred to a Geltrex-coated dish, incubated for 30 min, and fixed with 4% PFA. Duplicate wells were prepared for propagating NPCs into secondary and tertiary neurospheres by mechanically dissociating cells, re-plating them, and allowing them to grow for an additional 7 days.

### RT-PCR of NPC, neuron, astrocyte, and oligodendrocyte markers

GFP and self-tracing NPCs were plated on 6-well plates coated with Geltrex at 1 × 10^6^ cells per well. One hundred μg/mL of spinal cord homogenate from naïve or 8-week injured SCI animals was used to treat cells for 1 week, with regular media changes performed every other day. NPCs cultured in SFM were plated as controls. After 1 week of treatment, mRNA was extracted from the cells and synthesized into cDNA. RT-PCR was conducted using the TaqMan probes detailed in Supplementary Table 1. Samples were run in triplicates with values normalized to GAPDH.

### Whole-cell patch clamp recording

A coverslip was placed in a perfused chamber mounted on a Nikon E600FN microscope equipped with infrared differential interference contrast optics (IR-DIC) and an epifluorescence system that included a computer-controlled excitation wavelength switcher Lambda DG-4 (Sutter Instruments) and a high-resolution digital CCD 55 camera (ORCA ER C4742-95-12, Hamamatsu). Images were acquired using NIS Elements (AR3.10, Nikon) software and analyzed using ImageJ version 1.52 (Wayne Rasband, NIH).

The chamber was perfused at a rate of 0.5 ml/min with a solution containing (in mM): 140 NaCl, 5 KCl, 2 CaCl2, 1 MgCl2. 10 HEPES, 15 D-glucose and 3 Na-pyruvate, pH adjusted to 7.2 with NaOH (osmolarity ∼295 mOsm, measured with a freezing point Advanced Micro-Osmometer).

The cells were visualized and selected for patch clamp recording using IR-DIC and fluorescence for GFP as well as mCherry using appropriate filter cubes (EX480/40x, BS 505, EM510 for GFP and EX535/50x, BS 565, EM610/75 for mCherry).

The patch pipettes were pulled on a laser-based programmable pipette puller (model P-2000, Sutter Instruments) and filled with a solution containing (in mM): 130 K gluconate, 10 HEPES, 1 EGTA, 0.1 CaCl2, 2 MgATP, 5 NaCl and 0.3 Na-GTP (pH 7.2 adjusted with KOH, osmolarity ∼280 mOsm), which had resistances between 3 and 5 MΩ. The patch pipette solution was supplemented with 0.5 mM Lucifer Yellow (LY, dilithium salt, MW 457). LY fluorescence was recorded using the same filter used for GFP. Due to a much brighter LY signal in cells injected with LY during whole-cell recording, cell processes could be visualized and were easily recognizable after formaldehyde-fixation of the coverslips post-recording.

The cells were approached with a patch pipette using an MPC-385 Motorized Micromanipulator (Sutter Instruments). The pipette was pressurized to 10 mm Hg during approaching the cells, and a gigaseal (2-4 GΩ) was formed by releasing the pressure after establishing a close contact of the pipette tip with the cell membrane. A whole-cell mode of recording was established by mouth-applied brief negative pressure to rupture the membrane and establish a direct connection between the pipette interior and the cell cytoplasm. The gigaseal formation was monitored using the Membrane Test feature of pClamp10 software (Axon Instruments, Molecular Devices).

The signals, low-pass filtered at 10 kHz and sampled at 20 kHz, were recorded with a computer controlled MultiClamp 700B amplifier, digitized with Digidata 1440 and processed/analyzed using pClamp10 software (all from Molecular Devices). Both voltage clamp and current clamp recording (current clamp) were performed. Capacitance compensation and bridge balance were performed using the automatic functions of the MultiClamp 700B amplifier. To record Na+ and K+ currents, in voltage-clamp mode the cells were held at -80 mV, and voltage was stepwise changed in 10 mV 56 increments between -110 mV and +40 mV. For testing the ability of cells to generate action potentials, in current clamp mode, the membrane potential was held at around -70 mV, and 50 and 250 ms-long current pulses were applied to depolarize the membrane to action potential threshold levels, and up to 0 mV or higher when no action potentials were detected. To detect spontaneous synaptic activity, 2 minute-long continuous (“gap-free”) voltage clamp and current clamp recordings were performed in each cell, with membrane voltage being held at around -70 mV. All recordings were done at room temperature.

### Immunocytochemistry of neuroglial markers

Cells were fixed in their respective wells for 20 min at room temperature with an 8% PFA in 30% sucrose solution, added at a 1:1 ratio of media to fixative for a final solution of 4% PFA. Coverslips were washed three times with 1X PBS (10 min). For intracellular staining, cells were subjected to a solution of 0.1% Triton in PBS to permeabilize the cell membrane and placed on ice for 3 min. Blocking of non-specific antigen binding was achieved using a solution of 5% BSA for 1 h at room temperature. Primary antibodies for TUBB3, MAP2, and GFAP were diluted in blocking solution and applied to the coverslips for overnight incubation at 4°C. WGA and TTC antibodies were also used to enhance tracer signal.

### Clip contusion-compression SCI model

Adult female RNU rats aged 8 to 10 weeks-old were anaesthetized with 5% isoflurane for rapid induction carried by 1:1 NO2-O2. Isoflurane was reduced to 2% for maintenance. Animals were placed on heating pads to maintain constant body temperature. An additional gauze bolster was placed under the torso to achieve optimal access to the laminae and spinal cord during surgery. The upper backs of the rats were shaved and cleaned with 2 wipes each of 70% ethanol and betadine. To ensure post-operative analgesia, animals were administered buprenorphine (0.05 mg/kg) subcutaneously. Ten mL saline was also administered to ensure hydration. A No.15 blade was used to create a midline dorsal skin incision from the occipital protuberance to the prominent T2 spinous process. A second incision was made along the ligamentum nuchae to reach the laminae while minimizing bleeding. The muscle layers were retracted to expose the paraspinal muscles and the attachments of the spinal and deep muscles were laterally removed from the vertebrae to the medial edge of the articular facets by scraping with a scalpel. The ligamentum flavum was removed from the rostral end of C6 to the caudal end of C7 using the tip of the scalpel. The laminae were removed using angled offset bone nippers. To this point, the procedure is considered a C6/7 laminectomy, and this was also performed on sham animals. To create space for the extradural passage of the modified aneurysm clip, a blunted dissection hook was inserted in the space between the C6/7 spinal roots bilaterally. The bottom blade of the clip was passed along the ventral surface of the spinal cord. To generate the injury, the clip was applied quickly with a closing force of 23 g and the compression was sustained for 1 min. Puncture of the dura or spinal cord resulted in euthanasia and exclusion from the study. Following removal of the clip, the retractors were released. Muscles were closed using 3-0 nylon sutures along the midline and the skin was closed using Michel suture clips. Animals were removed from isoflurane and allowed to recover under a heat lamp in a clean cage with softened food and water on the floor.

### Cell-transplantation

Self-tracing NPCs and control GFP-NPCs were cultured in N2/B27 serum free media. Prior to transplantation, cells were collected in SFM at a density of 200 000 cells/μL. Cells were kept on ice until they were ready for use. Each sample could only be used for up to 2 animals (2 h max). Two weeks or 8 weeks post-injury, animals underwent similar surgical procedures as described above (see 2.5.1 Clip contusion-compression SCI model) with slight modifications. The scar tissue was carefully removed by working from the rostral and caudal ends of the C6/7 region. Using the tip of a 25G needle and a pair of fine forceps, the dura was opened. Animals were transplanted at 5 sites: directly in the epicentre cavity, and other 4 sites (1 mm rostral and caudal to the lesion at a depth of 1 mm, bilaterally). 2 μL of the cell suspension was infused into the epicentre while 1 μL was used in the penumbra. A 10 μL Hamilton syringe and microinjector pump was used to regulate the volume and rate of cell transplantation (1 μL/min). Injection needles were left to dwell for 2 minutes after injection to prevent backflow. The meniscus was checked before each transplant and the syringe was thoroughly flushed with saline between each animal to ensure there was no blockage.

### Immunohistochemistry (IHC)

Animals underwent anesthesia and transcardial perfusion at 2 weeks or 8 weeks post-SCI. The brain and spinal cord were removed from the animals and post-fixed with 10% sucrose in 4% PFA solution overnight. The following day, the tissue was washed with 0.1 M PBS and cryoprotected in 30% sucrose in PBS for 48 h. A section of the spinal cord measuring 2.5 cm in length surrounding the lesion epicentre was embedded in M1 embedding matrix and stored at -80°C. Brains were embedded in Tissue-Tec Optimal Cutting Temperature and stored at -80°C. Tissues were sectioned using a cryostat in 30 μm thick sections in either transverse or longitudinal sections. For immunohistochemistry, slides were thawed, dried, and rehydrated with 1X PBS. Nonspecific binding was prevented using a 1h incubation with blocking solution (5% milk, 1% BSA, 0.3% Triton) at room temperature. Primary antibodies for WGA and TTC were diluted in blocking solution and incubated overnight at 4°C. Three 15 min washes with PBS were performed before applying secondary antibodies with DAPI (1:1000) for 1 h at room temperature. Upon an additional three PBS washes, Mowiol was applied to the slides to mount coverslip. Slides were allowed to dry at room temperature overnight and transferred to 4°C for long-term storage.

### Statistics

Quantitative data are expressed as mean ± SEM. Equal variances between treatments, homogeneity of variances, and normal distribution of the data favoured the use of parametric statistical tests. Therefore, differences between groups were assessed by one- and two-way analysis of variance (ANOVA) with Tukey’s post hoc test to correct for multiple comparisons (p < 0.05). Data were analyzed with GraphPad Prism V6(GraphPad Software Inc., La Jolla, CA, US).

## Abbreviations

CMV: Cytomegalovirus
CNS: Central Nervous System
EF1a: Elongation factor 1 alpha
GAPDH: Glyceraldehyde 3-phosphate dehydrogenase
GFP: Green fluorescent protein
HRP: Horseradish Peroxidase
IPSC: Induced pluripotent stem cell
IRES: Internal ribosomal entry site
LY: Lucifer yellow
MAP2: Microtubule-associated protein 2
NIM: Neural Induction Media
NMM: Neural Maintenance Media
NPC: Neural progenitor cell
SCI: Spinal Cord Injury
pB: PiggyBac
TTC: Tetanus toxin fragment C
WGA: Wheat germ agglutinin

## Declarations

### Ethics approval and consent to participate

All animal experiments were approved by the animal care committee at the University Health Network (Toronto, Ontario, Canada) in compliance with the Canadian Council on Animal Care.

### Consent for publication

Not applicable.

### Availability of data and materials

The datasets used and/or analyzed during the current study are available from the corresponding author on reasonable request.

### Authors’ contributions

PC conducted experiments and performed cell culture, generation and differentiation of cells, cell characterization, quantitative PCR, in vitro studies, microscopy, and analysis of the data ZL and CSA assisted in cell culture and cell preparation animal surgeries, cell transplantation and histological analysis of the tissue. AV performed patch clamp and in vitro electrophysiology. MGF and MK conceptualized this study and established methodology. MGF provided resources. PC wrote the original draft. ZL, CSA, MK and MGF reviewed and edited the draft.

## Acknowledgments

We thank Dr. Tim Worden for copyediting the manuscript. We wish to recognize Jian Wang and Behzad Azad for their technical assistance and Sin Ki Kylie Lau, and Si Yu Chen for helping in animal care.

