## Supplemental Tables and Figures for "Development and Characterization of Self-Tracing Neural Progenitor Cells for Mapping Their Synaptic Integration into Endogenous Neural Networks"

**Supplemental Material**

**Supplementary Table 1 - RT-PCR Probe Catalogue Numbers from Applied Biosystems**

| Gene of Interest | TaqMan Probe |
| --- | --- |
| GAPDH | Hs02786624_g1 |
| NES | Hs04187831_g1 |
| SOX1 | Hs01057642_s1 |
| SOX2 | Hs01053049_s1 |
| DLG4 | Hs01555373_m1 |
| GAP43 | Hs00967138_m1 |
| TUBB3 | Hs00801390_s1 |
| AQP4 | Hs00242342_m1 |
| GFAP | Hs00909233_m1 |
| APC | Hs01568269_m1 |
| PDGFRA | Hs00998018_m1 |

**Self-tracing NPCs express typical NPC markers**

To validate that NPC phenotypes of self-tracing cells were reflected at the molecular level, RT-PCR was used to evaluate expression levels of known NPC markers in self-tracing NPCs post-FACS expansion against GFP NPCs (Supplementary Figure 1A). Generally, self-tracing NPCs showed similar expression levels in all markers to control cells. (one-way ANOVA, Tukey’s post hoc, p < 0.05). Moreover, ICC confirmed positive staining of nestin in self-tracing NPCs.

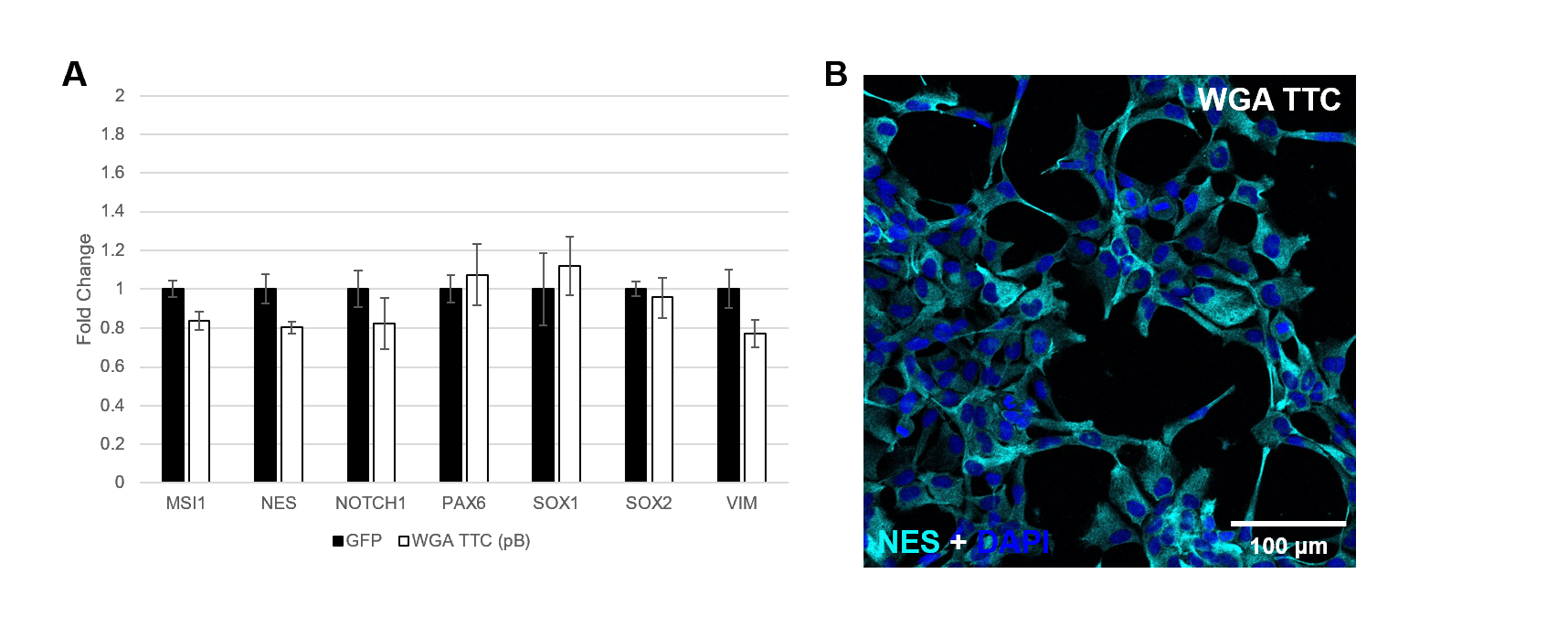

**Supplementary Figure 1: Self-tracing NPCs express typical NPC markers.** (A) GFP and both lines of genetically engineered self-tracing NPCs demonstrate no difference in NPC marker expression as assessed by RT-PCR. All groups were compared to wild-type human iPSC-derived NPCs. Error bars display SEM, n=5 per group. One-way ANOVA corrected for multiple comparisons by Tukey’s post hoc test, *p < 0.05. (B) Representative confocal image of self-tracing NPCs show expression of nestin. Abbreviations; NES = nestin; TTC = tetanus toxin fragment C; WGA = wheat germ agglutinin

**Profiling self-tracing NPC differentiation *in vitro* by RT-PCR**

Self-tracing NPCs were treated with either naïve or 8 week-injured spinal cord homogenates to induce differentiation of cells into their neuroglial fate over the course of 7 days. GFP NPCs were also used as controls, and relative expression changes were compared to SFM-cultured NPCs. Once mRNA was isolated and converted to cDNA, RT-PCR of NPC, neuron, astrocyte, and oligodendrocyte markers was performed (Supplementary Figure 2). Fold changes have been log2 transformed.

Compared to SFM-cultured NPCs, GFP and both lines of self-tracing NPCs demonstrated a slight increase in NPC markers, namely SOX1 and SOX2 (two-way ANOVA, Tukey’s post hoc, p < 0.001) when treated with naïve homogenate, but a decrease in expression with SCI 64 homogenate was observed. This suggests factors within the SCI microenvironment possibly promote differentiation of NPCs to neuroglial fates.

Upon naïve spinal cord homogenate treatment to NPCs, a slight increase in neuron marker expression was seen, which was most statistically significant for the marker DLG4 (GFP: 1.39 ± 0.26; WGA/TTC: 1.18 ± 0.19, p < 0.01), a gene involved in recruiting receptors, ion channels, and signaling proteins to post-synaptic sites. Similar patterns of upregulation in the naïve spinal cord microenvironment were seen with both astrocyte and oligodendrocyte markers. The greatest upregulation was seen with AQP4 (GFP: 11.70 ± 0.09; WGA/TTC: 11.07 ± 0.09, p < 0.0001), a gene encoding water channels, localized to astrocytic endfeet. Oligodendrocyte markers, such as APC, showed slightly less upregulation (GFP: 2.84 ± 0.27; WGA/TTC: 1.89 ± 0.08, p < 0.0001). These findings hint that NPCs are more likely to differentiate into astrocytes and oligodendrocytes than neurons.

After NPCs were treated with SCI homogenate for 1 week, a decrease in neuron markers was detected. Interestingly, GAP43 expression (GFP: -1.32 ± 0.12; WGA/TTC: -2.21 ± 0.11) was not only lower than in naïve conditions (GFP: 0.59 ± 0.42; WGA/TTC: - 0.28 ± 0.08, p < 0.0001), but also in control SFM conditions (GFP: N/A; WGA/TTC: -1.64 ± 0.14, p < 0.01). Astrocyte and oligodendrocyte markers were generally only slightly elevated compared to control conditions. Only AQP4 expression remained elevated (GFP: 7.65 ± 0.10; WGA/TTC: 6.73 ± 0.11, p < 0.0001). Astrocytes appear to be the predominant fate of NPCs in an SCI microenvironment.

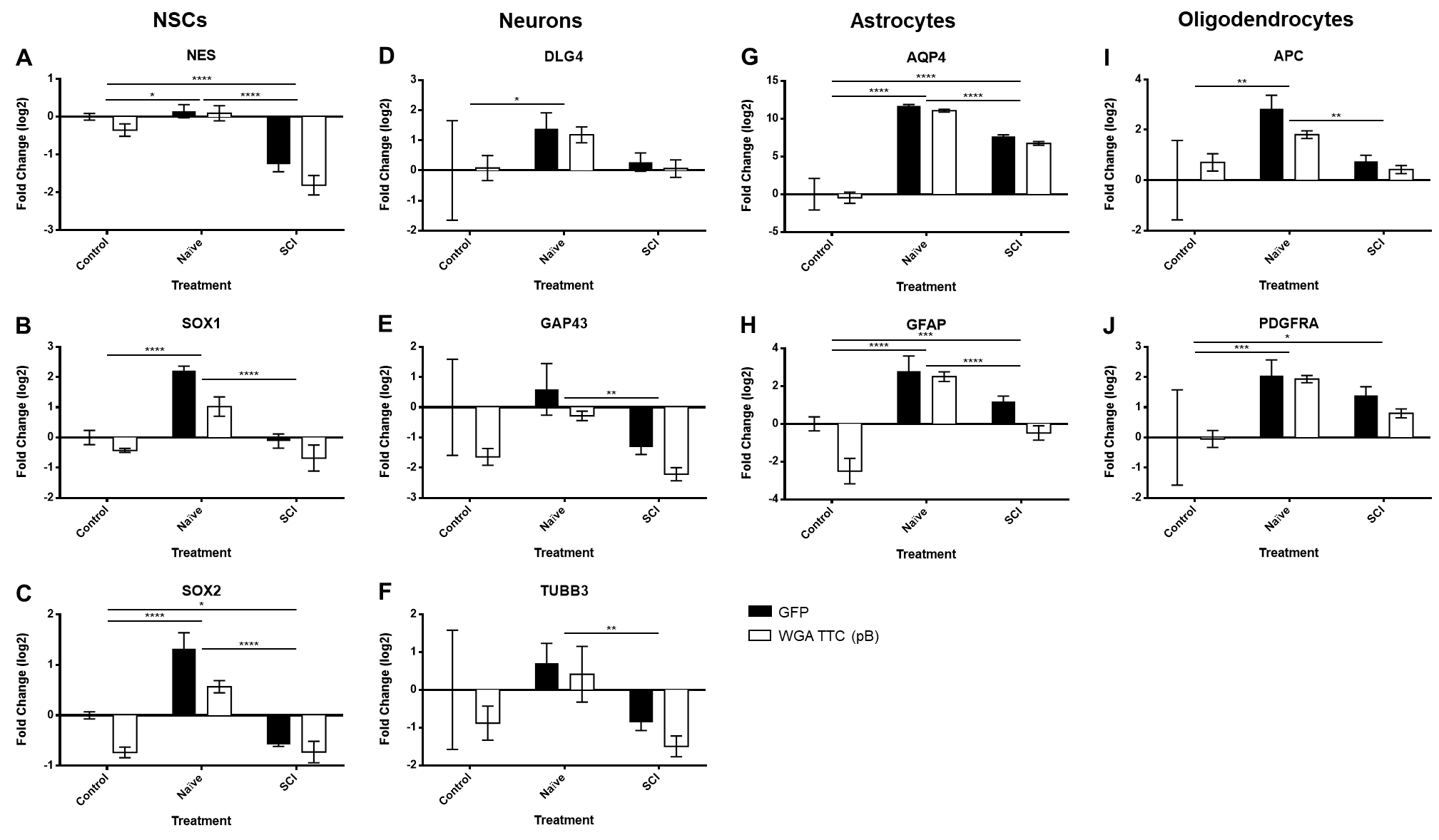

**Supplementary Figure 2: Differentiation profile of self-tracing NPCs by RT-PCR.** GFP and self-tracing NPCs were treated with spinal cord homogenate from a naïve or 8-week injured RNU rat for 1 week. NPCs cultured in SFM were used as a control. (A-C) NPC, (D-F) neuron, (G,H) astrocyte, and (I,J) oligodendrocyte markers were examined. Error bars display SEM, n=3 per group. Two-way ANOVA corrected for multiple comparisons by Tukey’s post hoc test, *p < 0.05; **p < 0.01; ***p < 0.001; ****p < 0.0001.
